# Immune Composition of the Mononuclear Cell Fraction of Human Umbilical Cord Blood

**DOI:** 10.1101/2025.04.21.649733

**Authors:** Karen Kikuta, Esmond Lee, Talia Menezes, Hannah Fung, Alvaro Amorin, Aditi Agrawal, Theodore L. Roth, Matthew Porteus

**Affiliations:** Center for Definitive and Curative Medicine, Stanford University School of Medicine, Stanford, USA; Department of Biology, School of Humanities and Sciences, Stanford University, Stanford, USA; Department of Pathology, Stanford University School of Medicine, Stanford, USA; Division of Pediatric Hematology, Oncology, and Stem Cell Transplantation and Regenerative Medicine, Stanford University School of Medicine, Stanford, USA; Institute for Stem Cell Biology and Regenerative Medicine, Stanford University School of Medicine, Stanford, US; Department of Pathology, School of Medicine, Stanford, USA; Chan Zuckerberg BioHub, San Francisco, USA; Bioprocessing Technology Institute (A*STAR), Singapore

## Abstract

Despite its therapeutic potential and unique immunological properties, the immune composition of umbilical cord blood lacks consistent and comprehensive characterizations. Human umbilical cord blood (UCB) is often discarded after delivery and is difficult to obtain for research purposes. Furthermore, most research on UCB is focused on properties of CD34+ hematopoietic stem cells for transplantation. The Binns Program for Cord Blood Research at Stanford University has the unique advantage of regular collection and isolation of mononuclear cells (MNC) from UCB donors. This study provides a robust characterization of the immune subset compositions of the CD34-negative MNC fraction of UCB (n=50). The study also compares the UCB data to adult peripheral blood (PB) mononuclear cells to identify differences in immune maturity. Using flow cytometry and single-cell RNA sequencing (scRNA-Seq), we analyzed UCB and adult PB MNC samples to characterize the cell surface protein and transcriptomic profiles of different immune subsets. Our study findings bring a higher-definition understanding of the unique immunological properties of umbilical cord blood. Study findings reveal a distinct immune profile in UCB, such as a higher average percentage of CD19 B Lymphocytes, CD4 T Cells, CD4 Naive T Cells, CD4 Recent Thymic Emigrants, CD8 Naive T Cells, CD8 Recent Thymic Emigrants, and CD19 Naive B Cells compared to adult PB. Additionally, there were fewer CD19 Memory B Cells in UCB compared to PB. The scRNA-Seq showed concordance in the proportion of immune cell types but captured more differentiated subtypes of cells. Additionally, scRNA-Seq showed unique clustering patterns in UCB, which reflect cell types that converge in adulthood as the immune system matures. These analyses yield the intriguing possibility that the immune heterogeneity of individuals at birth gives way to more stereotyped immune subsets as the immune system is exposed to the external environment and undergoes maturation. Overall, our findings provide a robust characterization of MNC UCB immune subsets and insights into how immune function develops from birth to adulthood.

## Introduction

Human Peripheral Blood Mononuclear cells (PBMCs) contain immune cells essential to innate and adaptive immunity (23). While conducting immune surveillance in systemic circulation, cells can also traffic to secondary lymphoid organs and tissues in response to infection and inflammation. Immune challenges over the lifetime of an individual modifies the immune system and leads to mature cells with immune memory (24). One way to understand a mature immune system is to understand its point of origin at birth. Mononuclear cells (MNCs) from Umbilical Cord Blood (UCB) provide the opportunity to study the naive immune system. A few studies have investigated the properties of UCB MNCs, typically focusing on the hematopoietic stem and progenitor cell (HSPC) population (4, 11, 5, 6, 7, 8) and making comparisons with bone marrow (1, 2, 3) derived HSPCs, which have been the gold standard in transplantation.

From an immunological perspective, early studies investigating the phenotype of UCB MNCs identified populations such as T- and B-lymphocytes, as well as NK cells (5). They showed that lymphocytes appeared to be phenotypically immature (6, 15). Notably, these studies were published over 20 years ago before technological advances allowed high-dimensional analysis of component cell populations. In recent years, single-cell transcriptomics was used to analyze the expression patterns of known marker genes of nucleated cord blood cells (21). Despite recent publications and the continued recognition of the promise of UCB research (3), comprehensive characterizations of the MNC populations of UCB are still lacking.

The Binns Program for Cord Blood Research gave us the opportunity of obtaining a large number of human UCB on a weekly basis (10). Annually, the program collects hundreds of UCB samples, and its established MNC isolation protocol has led to the distribution of UCB products to over 20 laboratories and over 60 researchers around the Stanford University campus. We studied the immune subsets found in 50 UCB donors by flow cytometry and paired this with single-cell transcriptomics (scRNA-Seq) to gain deeper insight into the complexity of the MNC fraction. Additionally, we compared the immune subtype composition of UCB to adult peripheral blood (PB) MNCs to understand changes that take place in immune maturity.

## Materials and Methods

### Study Design

Immunophenotyping of mononuclear cells (MNCs) using flow cytometry was performed on 50 umbilical cord blood (UCB) and 22 adult peripheral blood (PB) samples. Additional analysis with 3 UCB and 3 PB samples was performed using RNA-sequencing for a more detailed characterization. Similar studies performed by D’Arena G et al. (1998) and Mantri S. et al. (2020), as well as resource availability (laboratory space, finances, and time), were used as a reference to determine sample size.

### Isolation of Mononuclear Cells

UCB MNCs were isolated from the UCB of term deliveries (≥34 weeks of gestation) at Lucile Packard Children’s Hospital-Stanford. UCB collections were performed through the Binns Program for Cord Blood Research with donor consent and institutional review board approval. Within 24 hours of collection, MNCs of UCB were obtained by density gradient separation of whole blood (Ficoll Paque Plus, GE Healthcare; 400g, room temperature, 30 minutes, deceleration off), followed by ammonium chloride red blood cell lysis (9:1 NH4Cl lysis buffer to cell suspension). Some UCB samples underwent additional processing to isolate CD34-negative MNCs by labeling with the human CD34 Microbead Kit Ultrapure according to manufacturer protocol (Miltenyi Biotec, San Diego, CA, USA) (10). Adult peripheral blood (PB) was collected from the Stanford Blood Center (SBC). Within 24 hours of collection, whole blood was diluted by 1:1 ratio with MACs buffer (1x PBS, EDTA, FBS). MNCs of PB were then obtained by density gradient separation (Ficoll Paque Plus, GE Healthcare; 400g, room temperature, 30 minutes, deceleration off), followed by ammonium chloride red blood cell lysis (9:1 NH4Cl lysis buffer to cell suspension). While our initial aims were to capture mononuclear cells, multinucleated cells such as Neutrophils found in the MNC fraction were also included as part of the study.

### Immunophenotyping of CD34-Negative Mononuclear Cells with Flow Cytometry

Following MNC isolation, 1 million cells per sample were stained with antibodies against markers at optimal concentrations according to manufacturer protocol (Supplemental Table 1). At least 100,000 events were acquired on a Beckman Coulter CytoFLEX flow cytometer and analyzed using FlowJo software (version 10.8.1) (Supplemental Figure 1). The study began with two panels to capture broad immune MNC subsets. They were updated to include new markers and a third panel to identify naive B cells (IgD), monocytes and granulocytes (CD66b), and TR1 cells (CD19b, LAG3). While our flow dataset has a total of 22 PB donors and 50 UCB donors, figures 2D, 2E, 3H, 4A & 4B, have 14 PB donors and 24 UCB donors, reflecting the donors sampled after the panels were updated.

**Figure 1.**
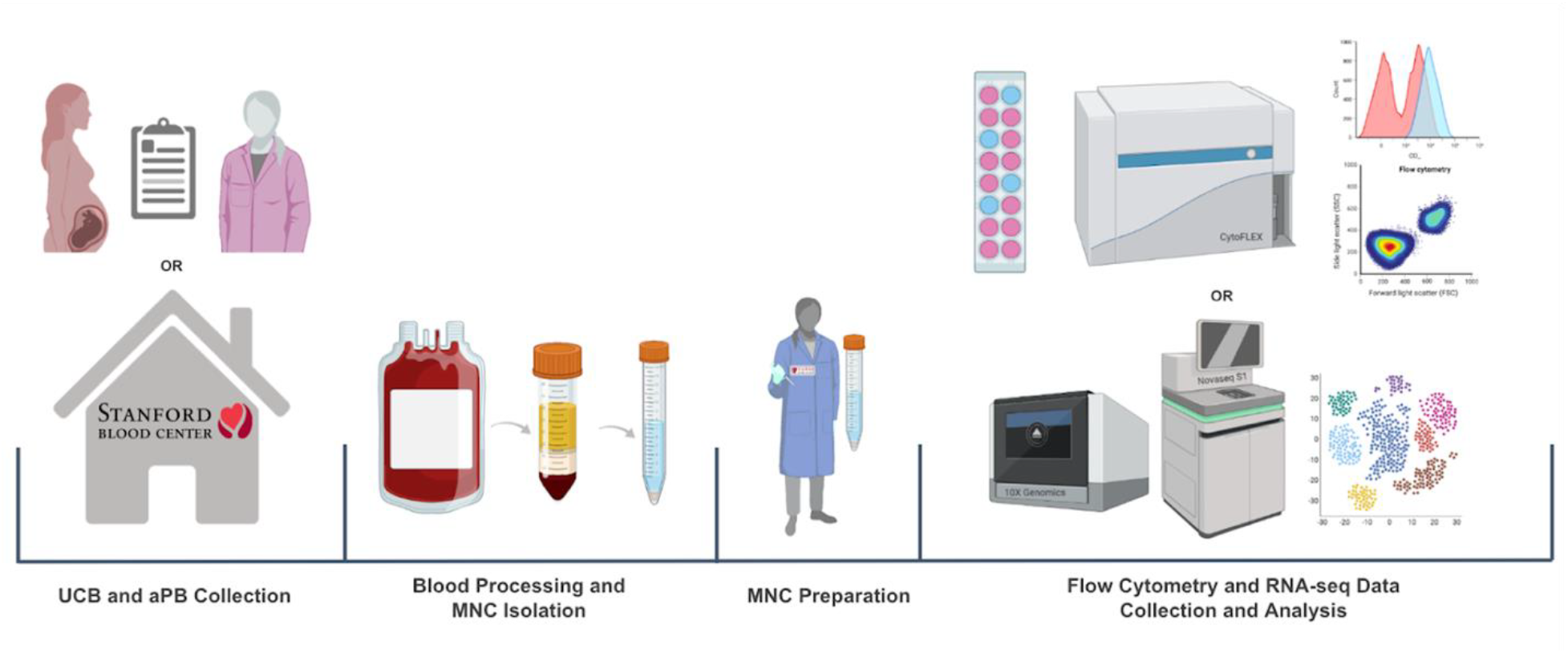
Umbilical Cord Blood (UCB) and Adult Peripheral Blood (PB) Workflow

**Figure 2.**
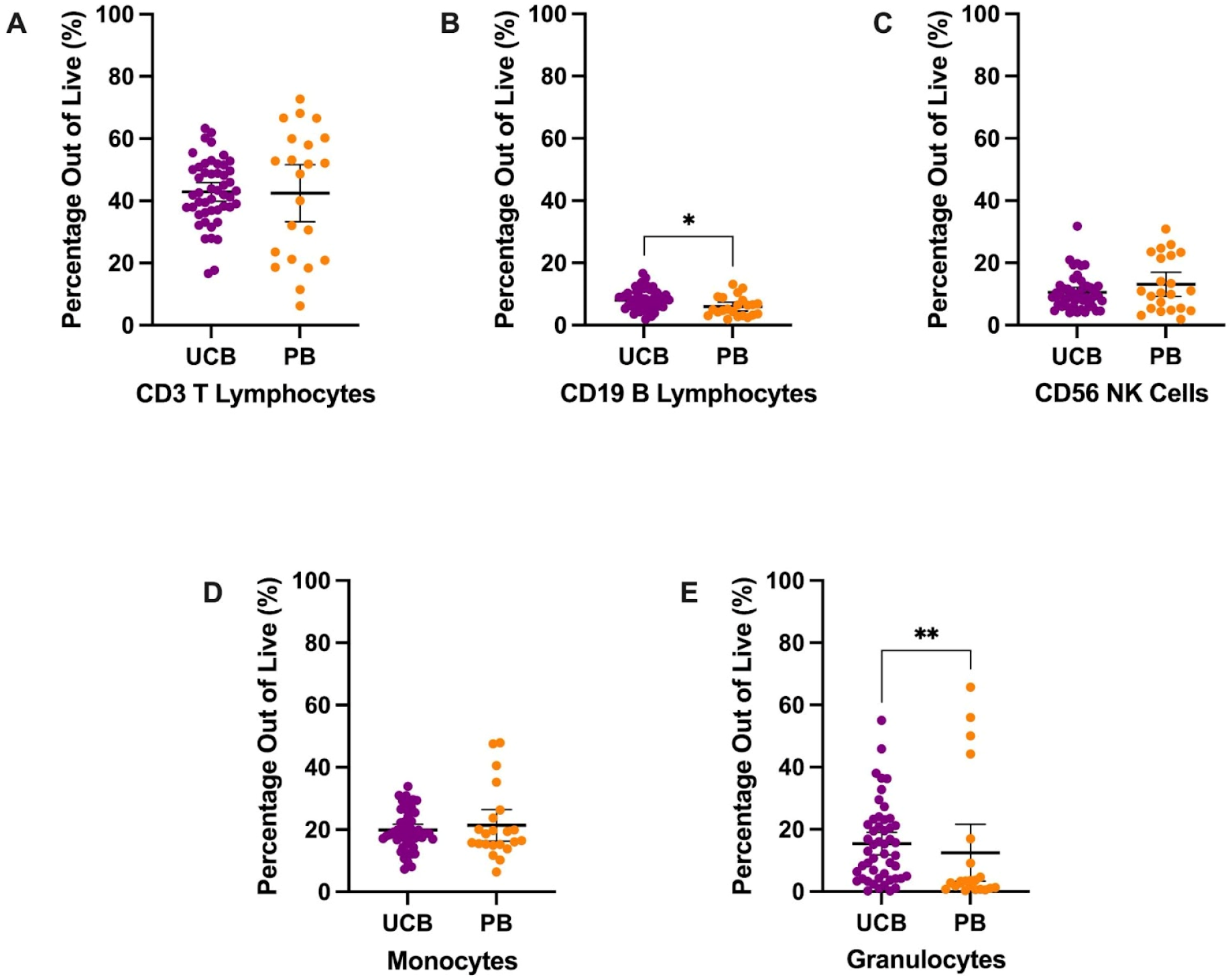
Overall Immune Composition by Mean with error bars reflecting the 95% Confidence Interval.

### Single Cell RNA-seq library preparation and sequencing

Libraries were prepared using Chromium Next GEM Single Cell 3′ Reagent Kits v3.1 single index kit according to the manufacturer’s protocol (10x Genomics), targeting 10,000 cells per sample. 12 cycles of cDNA amplification were done for all samples. Individual libraries were quality checked on an Agilent 4200 Tapestation 827 using D5000 screen tape. Next, KAPA library quantification kit (#KK4923) was used for qPCR on a BioRad CFX96 RT PCR thermal cycler. Single index libraries were sequenced on.

### QC of RNA-seq data

10x Genomics Cell Ranger was used to filter and align FASTQ files. Data analysis was performed in an R environment with Seurat (27) (https://satijalab.org/seurat/). Data were first filtered for quality based on number of unique features as well as mitochondrial counts. QC criteria were 200<nFeature<95th percentile and % Mitochondrial reads<98th percentile per cell for each donor. Log-normalization was performed on each cell to normalize feature expression measurements by total expression. These were performed according to donor type (UCB or PB) separately (Supplemental Table 4). The two Seurat objects were then merged for scaling and clustering. (Fig 5B). Cell clusters were labeled based on lineage specifying genes (Supplemental Table 5) and clusters that were attributed to a single donor were specified.

### Clustering and labelling of RNA-seq data

Clustering in Seurat is based on finding a subset of features exhibiting high variability in expression in the dataset. These were identified using the ‘VariableFeatures’ function. The counts were then scaled before performing dimensional reduction using Principal Component Analysis (PCA). We then used Uniform Manifold Approximation and Projection (UMAP) to project cells in two dimensions, with similar cells being plotted closer together. Known hematopoietic lineage markers were projected onto cell populations using ‘FeaturePlot’ to identify hematopoietic and immune cell types that could be compared with flow cytometry data. Elbow plots reflecting the standard deviations of the principle components were used to identify the number of significant dimensions that were used for clustering.

### Statistics

Statistical analysis was performed by GraphPad Prism (Version 9.4.0) and SAS® Studio (Release 3.81; Enterprise Edition). Data reported in the figures reflect the mean with a 95% Confidence Interval and we report mean and Standard Deviation (SD) in the text, assessing the heterogeneity within UCB MNCs samples and their specific immune subtypes. The relationship between UCB and PB MNCs was assessed using a parametric unpaired t-test with Welch’s correction or a nonparametric Mann-Whitney test based on Shapiro-Wilk normality test results. Detailed statistical results can be found in Supplemental Tables 2 and 3.

## Results

### Umbilical Cord Blood Results

UCB was obtained through the collaboration between the Binns Program for Cord Blood Research and Lucile Packard Children’s Hospital. PB from healthy adults was obtained through the Stanford Blood Center. The collected whole blood samples were then processed through density gradient separation to isolate MNCs. For flow cytometry, MNCs were stained with antibodies against immunologic markers of interest, and data was collected with CytoFLEX and analyzed with FlowJo. For RNA-seq, the samples went through library prep (10x genomics), QC, sequencing (Novaseq S1), and analyzed with Seurat.

### Flow Cytometry Immune Cell Population Characterization

MNCs were stained and analyzed to describe the populations’ major immune subpopulations, including T lymphocytes, monocytes, granulocytes, natural killer (NK) cells, and B lymphocytes (Fig 2). The largest mean proportion of MNCs in UCB in Panel 1 was identified as T lymphocytes (CD3+) at an average of 42.89% (SD: 10.37) of the live cells (Fig 2A). The smallest proportion of MNCs in UCB was, on average, 8.07% (SD: 3.17) B lymphocytes (CD19+) (Fig 2B). UCB composition was, on average, 19.85% monocytes (SD: 6.38) and 10.57% NK cells (SD: 5.44) (Fig 2C & 2D). Additionally, we found UCB to have an average of 15.40% granulocytes (SD: 12.67) (Fig 2E). The remainder of the UCB and PB descriptive statistics can be found in Supplemental Table 2. The immune composition of UCB compared to PB did not significantly differ for T lymphocytes, NK cells, monocytes, and granulocytes (p>0.05). However, UCB had a 2.13% higher mean proportion of B lymphocytes compared to PB (p=0.0125) (Supp Table C).

CD3+ T lymphocyte populations (Fig 3) make up a large percentage of the live mononuclear cells seen in umbilical cord blood and adult peripheral blood. In UCB and PB, the T lymphocyte population is mostly composed of CD4 T cells and CD8 T cells (Fig 3). On average, the T lymphocyte cells of UCB were 71.95% CD4 T cells (SD: 6.03) and 25.33% CD8 T cells (SD: 6.04) (Fig 3A & 3B). UCB had a mean CD4 T cell proportion that was 6.73% higher than the proportion for PB (p=0.0265) (Supplemental Table 3). There was no significant difference in CD8 T cells between UCB and PB (p>0.05) (Supplemental Table 3).

**Figure 3.**
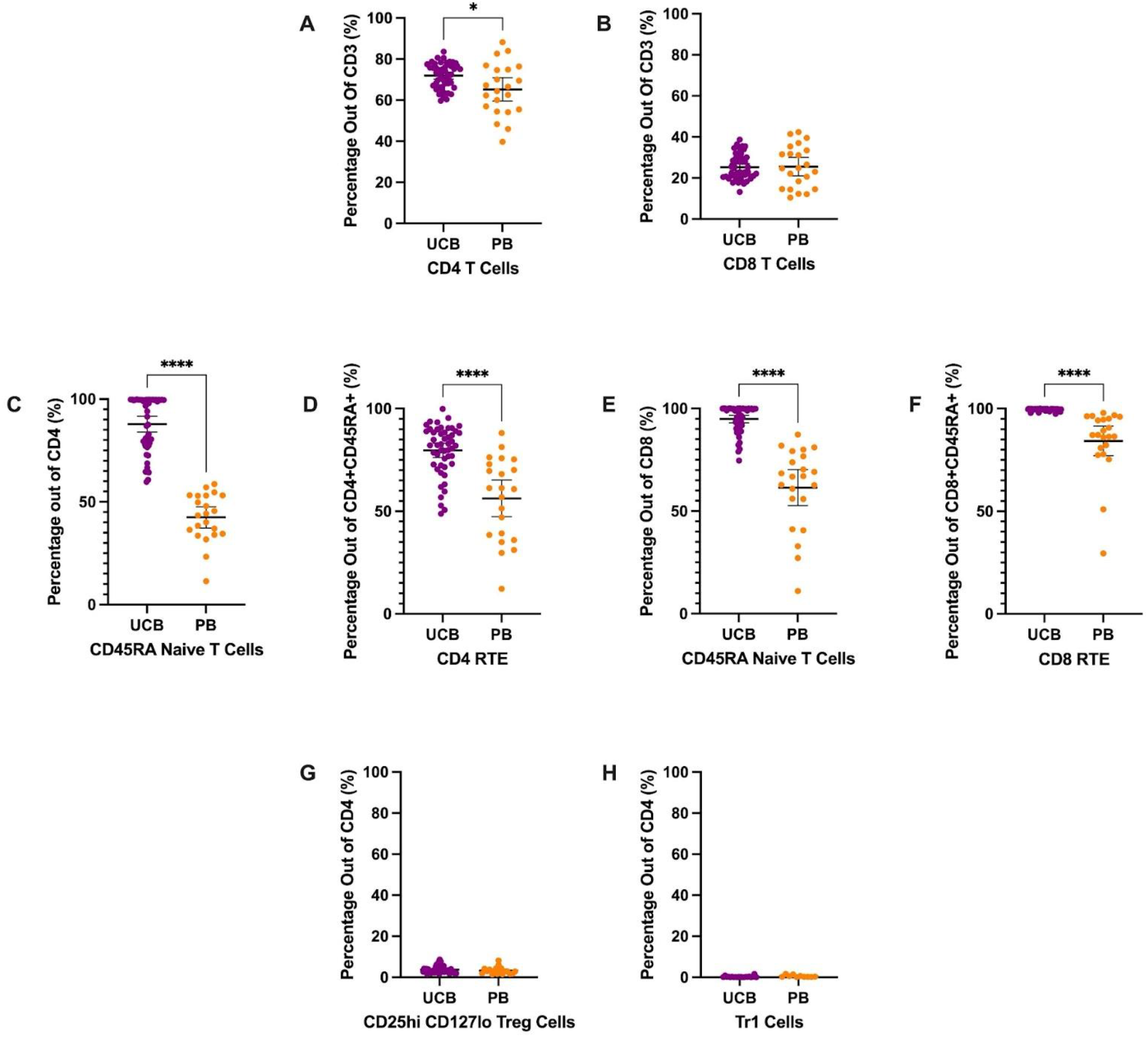
T Cell Subtype Composition by Mean with error bars reflecting the 95% Confidence Interval.

Looking closer at the subtypes of CD4 T cells, a large proportion were CD4 CD45RA naive T cells in UCB. Out of CD4 T cells, 87.79% were CD4 naive T cells (SD: 13.35), (Fig 3C). UCB had a mean CD4 naive T cell count that was double the mean count in PB (p<0.0001) (Supplemental Table 3). Within the CD4+ naive T cell population, there was also a significantly higher average cell count of recent thymic emigrants (RTE) cells in UCB compared to PB (p<0.0001) (Supplemental Table 3; Fig 3D). UCB had an average of 79.67% RTE subpopulation (SD: 12.14) within the CD4 CD45RA naive T cells. Comparatively, the mean percentage of RTE cells within the CD4 naive T cells in the PB samples was 23.39% lower (Supplemental Table 3).

From the CD4 T cells, there were also small subpopulations of Regulatory T (Treg) and Type 1 regulatory cells (TR1) in both UCB and PB (Fig 3G & 3H). The UCB samples had an average of 3.71% (SD: 1.69) and 0.27% (SD: 0.37) from the CD4 T cell population identified as Treg (Fig 3G) and TR1 cells (Fig 3H), respectively. No significant difference was found in the mean subpopulation percentages for TR1 or Treg cells between UCB and PB (p>0.20) (Supplemental Table 3).

When analyzing the subpopulations of CD8 T cells, the majority were CD8 CD45RA naive T cells in UCB and PB (Fig3E). On average, CD8 T cells in UCB had 94.87% (SD: 6.52) cells identified as CD45RA naive T cells. Similar to the CD4 naive T cells, UCB had a 33.39% higher proportion of the CD8 T cells as naive compared to PB (p<0.0001) (Supplemental Table 3). Of these CD8 CD45RA naive T cells, an average of 99.53% (SD: 0.55) were recent thymic emigrants (RTE) in UCB. There was a 15.23% higher proportion of RTE cells in UCB CD8 naive T cells compared to PB CD8 naive T cells (p=0.0002) (Supplemental Table 3).

Mononuclear cells were also stained and analyzed to characterize natural killer (NK) cells and B cells (Fig 4). Mature NK cells were gated from the CD3-CD19-CD56+ population (Supplemental Figure 1) (6, 14). On average, 73.32% of the CD56+ cells (SD: 15.55) were mature NK cells in the UCB samples (Fig 4C). The difference in percentage of mature NK cells between UCB and PB samples was not significant (p>0.05) (Supplemental Table 3).

**Figure 4.**
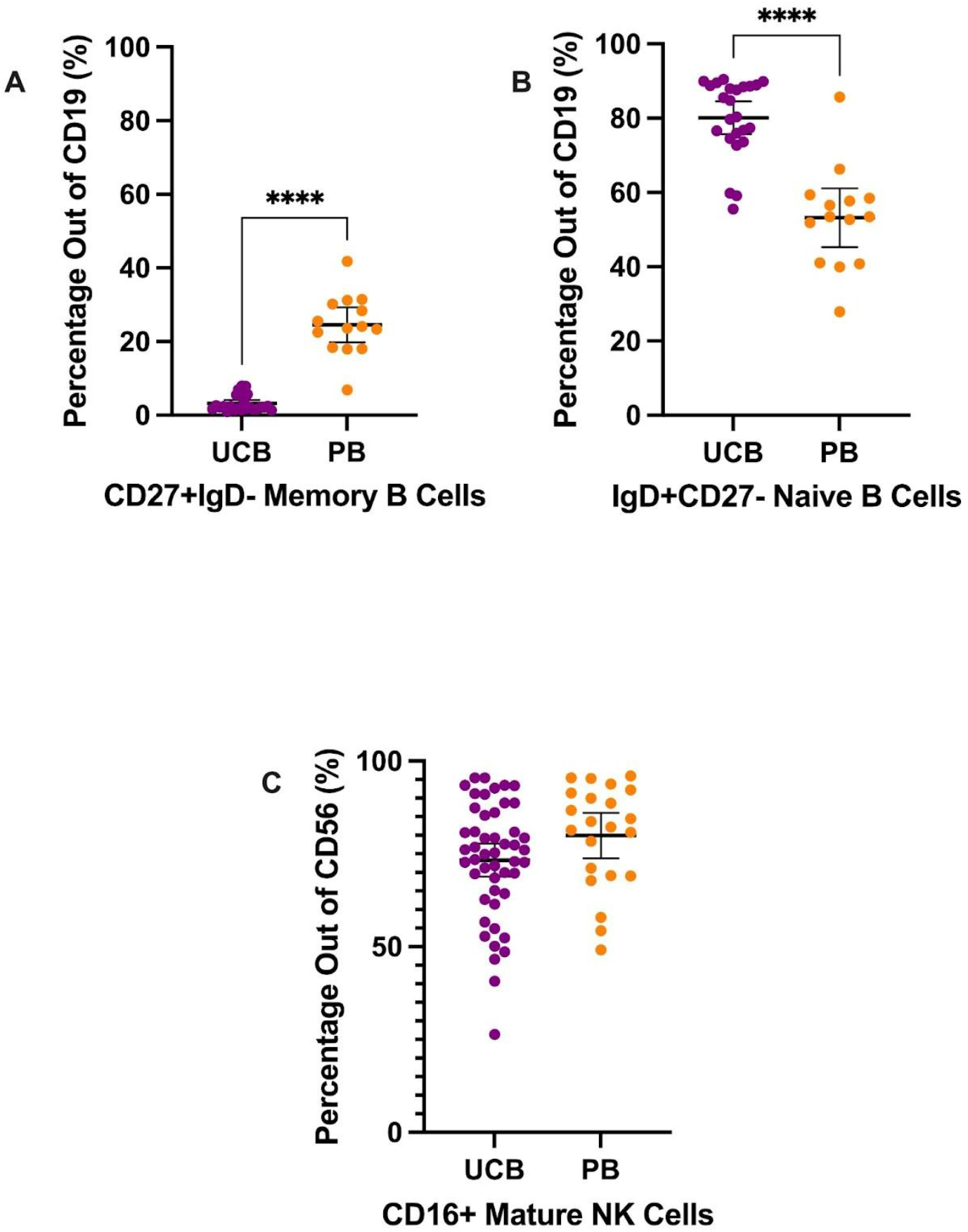
B Cell and NK Cell Subtypes Composition by Mean. Error bars reflect the 95% Confidence Interval

**Figure 5.**
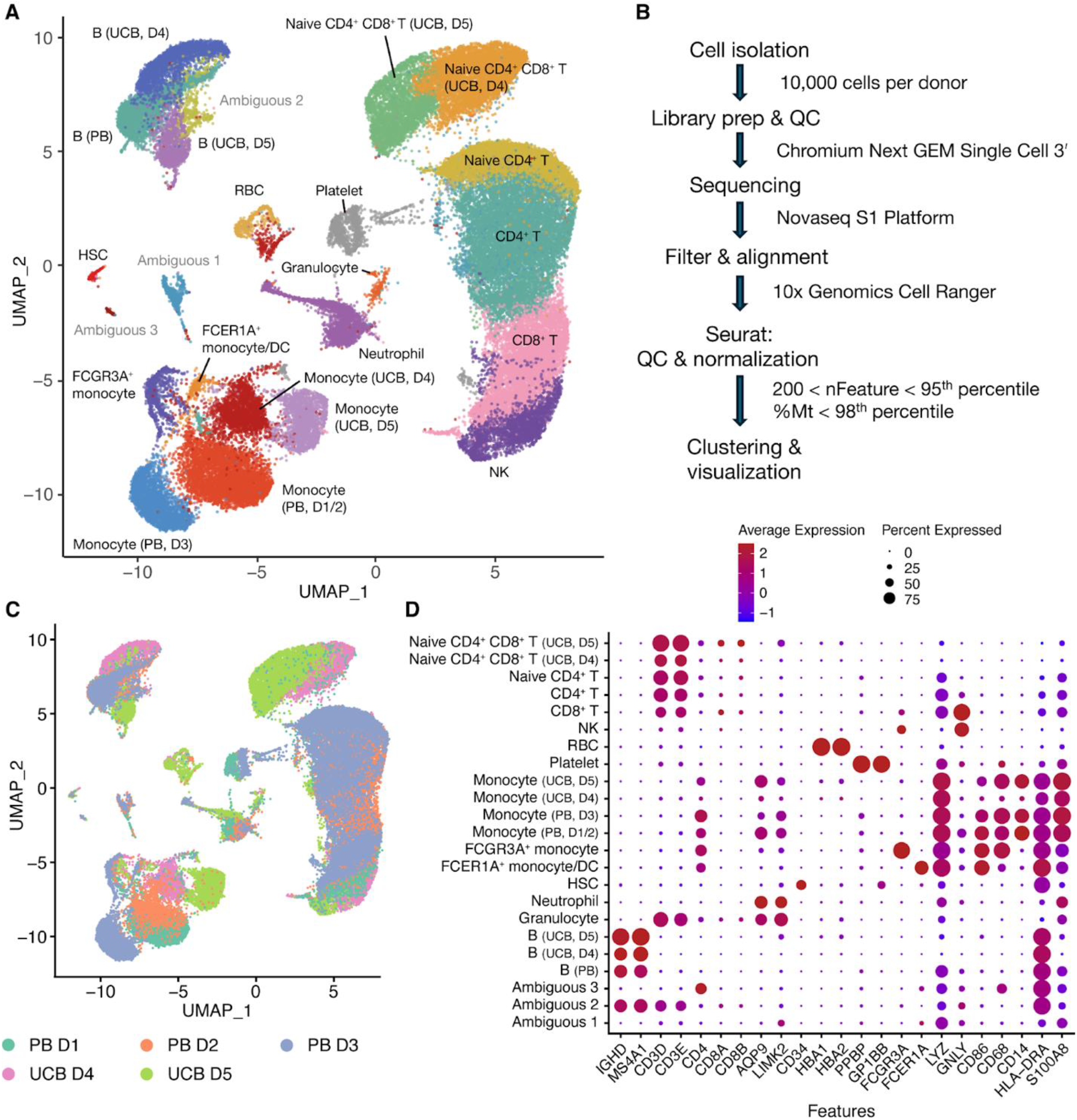
: RNA-seq data showing cell clusters from UCB and PB donors (A) UMAP plot of clustered PB (n=3) and UCB (n=2) donor cells. D stands for donor. (B) Flow chart of analysis (C) UMAP plot of clustered PB and UCB donor cells by donor contribution (D) Dotplot of cell clusters based on lineage-specifying genes.

Lastly, we analyzed the mature vs naive subtypes of B lymphocytes (Fig 4A & 4B). From the CD19+ B lymphocyte population, UCB had a dramatically larger mean proportion of naive B cells compared to its mean proportion of memory B cells—80.12% (SD: 10.40) versus 3.19% (SD: 2.23), respectively (Fig 4B & 4A). Meanwhile, the B lymphocytes in PB samples were, on average, 53.24% naive B cells (SD: 13.71) and 24.54% memory B cells (SD: 8.22) (Fig 4B & 4A). From their B lymphocyte populations, UCB had a mean proportion of memory B cells nearly eight times smaller compared to PB (p<0.0001) (Supplemental Table 3). Additionally, the proportion of naive B cells from the B lymphocytes was an average of 26.88% higher in UCB than in PB (p<0.0001) (Supplemental Table 3).

### sc-RNAseq Results

#### Combined UCB-PB scRNA-seq clustering

After library prep and sequencing, read alignment was performed with 10x Genomics Cell Ranger 7.0.1 using 10x Genomics Cloud Analysis (Fig 5B). One UCB donor was excluded from analysis because Cell Ranger detected an estimated number of 111,851 cells when only 10,000 cells were used for library prep. For the donor samples included in the analysis, we proceeded to data processing using Seurat. Cells were filtered based on having between 200 and the 95th percentile of nFeatures, as well as less than the 98th percentile of percentage mitochondrial reads per donor (Fig 5B). After QC, an average of 92.8±0.4% of cells were retained (Supplemental Table 4) across all donor samples. We obtained 36,915 cells from PB donors (n=3 donors) and 21,728 cells from UCB donors (n=2 donors). The normalized UCB and PB scRNA-seq datasets were merged to create a combined dataset of 58,643 cells (Fig 5A). Clustering analysis separated cells into 25 clusters which were annotated based on lineage-specifying genes (Fig 5A,C,D and Supplemental Figure 2) from the literature (24, 27, 29, 30). Clusters without a clear cell lineage were labeled ambiguous.

#### Concordance between flow cytometry and scRNA-seq data

In addition to analyzing the merged datasets, we clustered the UCB and PB data separately resulting in 15 clusters for PB donors (Fig 6A) and 20 clusters for UCB donors (Fig 6B). These were annotated based on lineage-specifying genes (Supplemental Figure 3 and Supplemental Table 5); clusters without a clear cell lineage were labeled ambiguous. The cell type proportions derived from our scRNA-seq analysis for the two UCB donors aligned well with the proportions derived from our flow cytometry data (Fig 6D, Table 2). Cell identities were obtained from flow through the markers CD13 (Monocytic), CD56 (NK), CD19 (B cell) and CD3 (T lymphocyte) from Panel 1. The proportion of CD4+ and CD8+ cells out of total CD3+ cells were obtained from Panel 2 out of total CD3 (Fig 6C). The data from flow cytometry contained more cells with an unknown identity because of the limited number of markers we could use to classify them. Both datasets showed a smaller proportion of B and NK cells, but a larger proportion of Granulocytes in UCB D5 compared to UCB D4. For example, UCB donor 5 had a smaller proportion of NK cells in both datasets (5.6% by flow, 7.1% by scRNA-seq) compared to UCB donor 4 (16.2% by flow, 16.7% by scRNA-seq) (Fig 6D, Table 2). The proportions of CD4, CD8 and NK cells were most similar across datasets. Using the Chi-squared test, we did not detect a significant difference in the cell type proportions identified by flow and scRNA-seq in both UCB donor 4 (χ2=0.152, df=6, p=0.999) and UCB donor 5 (χ2=0.128, df=6, p=0.999).

**Table 1.**
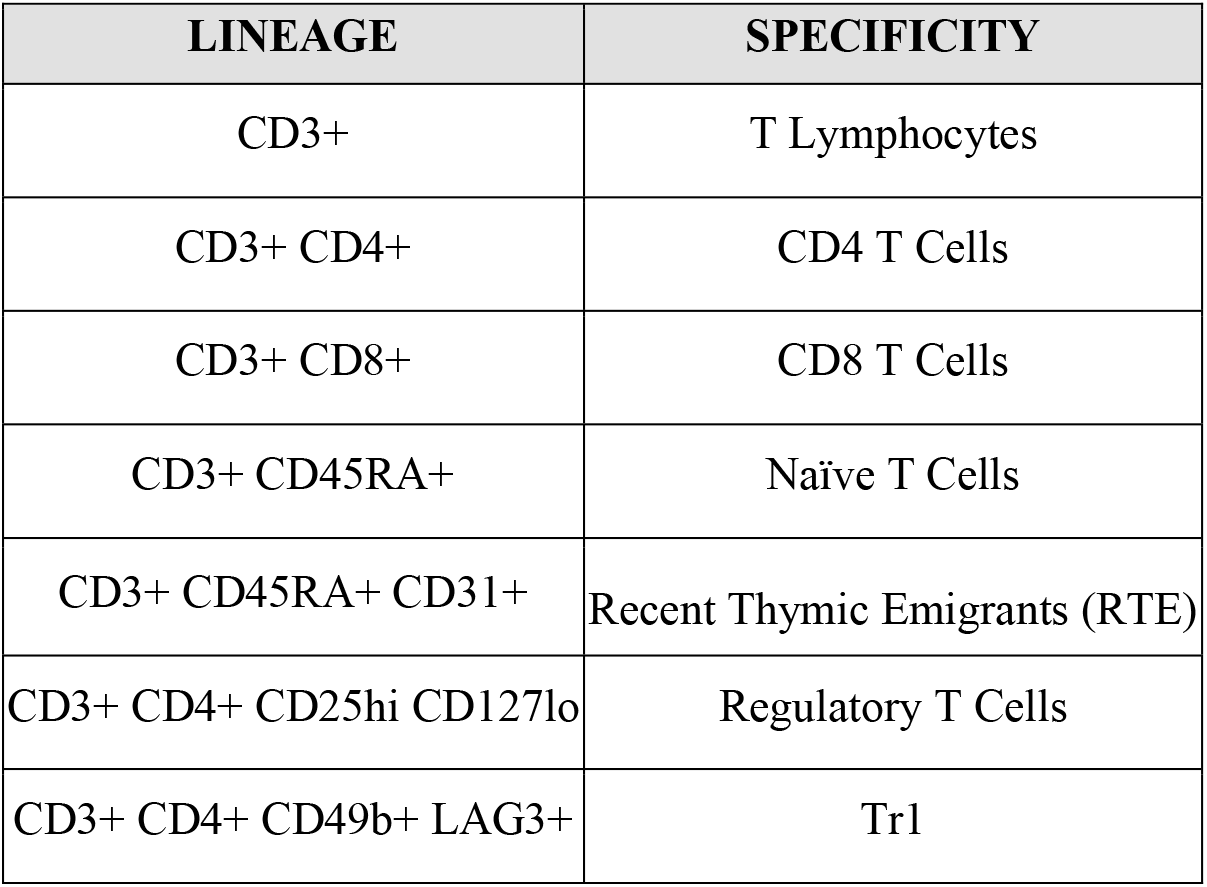
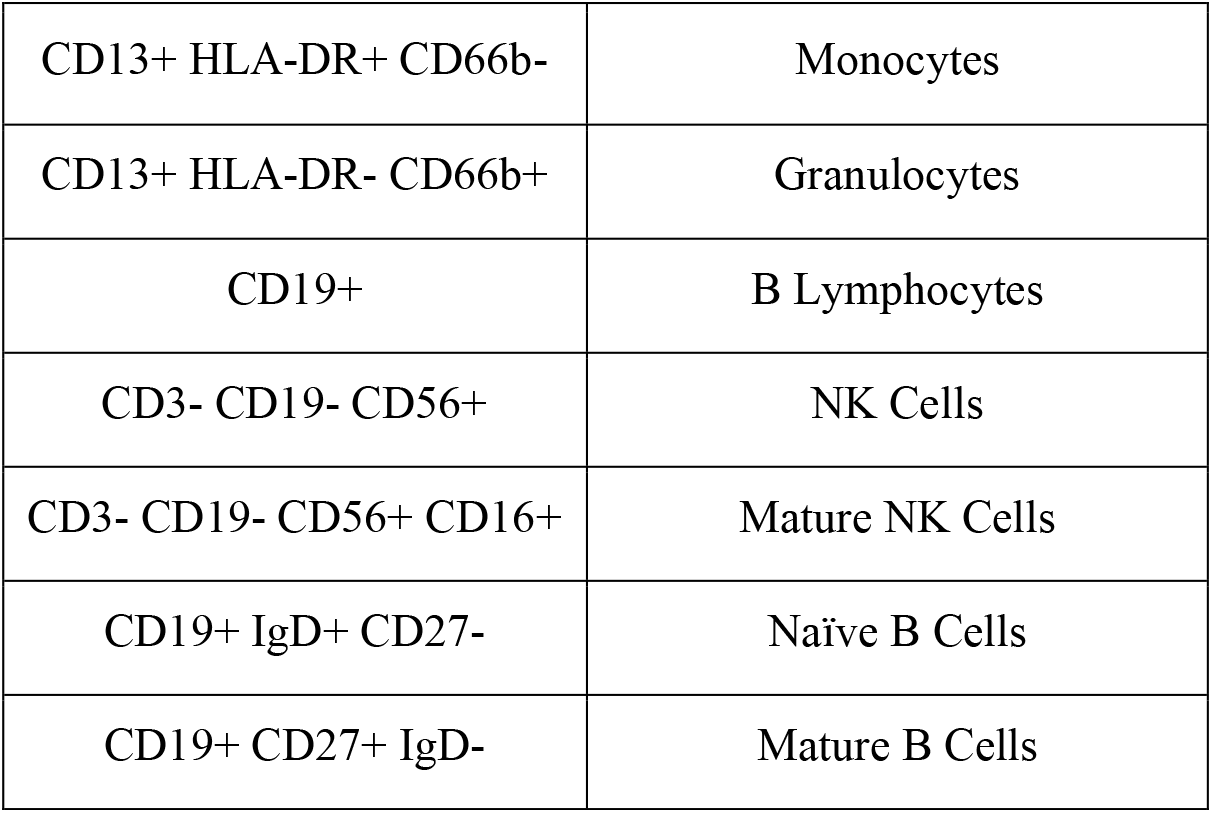
Cell Population and Specificity.

**Table 2.** Cell-type proportions from UCB MNC data from flow cytometry and scRNA-seq.

|  | CD4 | CD8 | Granulocyte | Monocytic | B cell | NK | Unknown |
| --- | --- | --- | --- | --- | --- | --- | --- |
| UCB D4 flow | 29.5% | 10.7% | 6.0% | 18.4% | 9.5% | 16.2% | 9.6% |
| UCB D4 RNA-seq | 27.2% | 10.5% | 0.0% | 19.6% | 23.2% | 16.7% | 2.8% |
| UCB D5 flow | 31.9% | 10.4% | 21.6% | 16.7% | 5.2% | 5.6% | 8.6% |
| UCB D5 RNA-seq | 32.2% | 12.6% | 11.4% | 21.8% | 13.1% | 7.1% | 1.8% |

**Figure 6.**
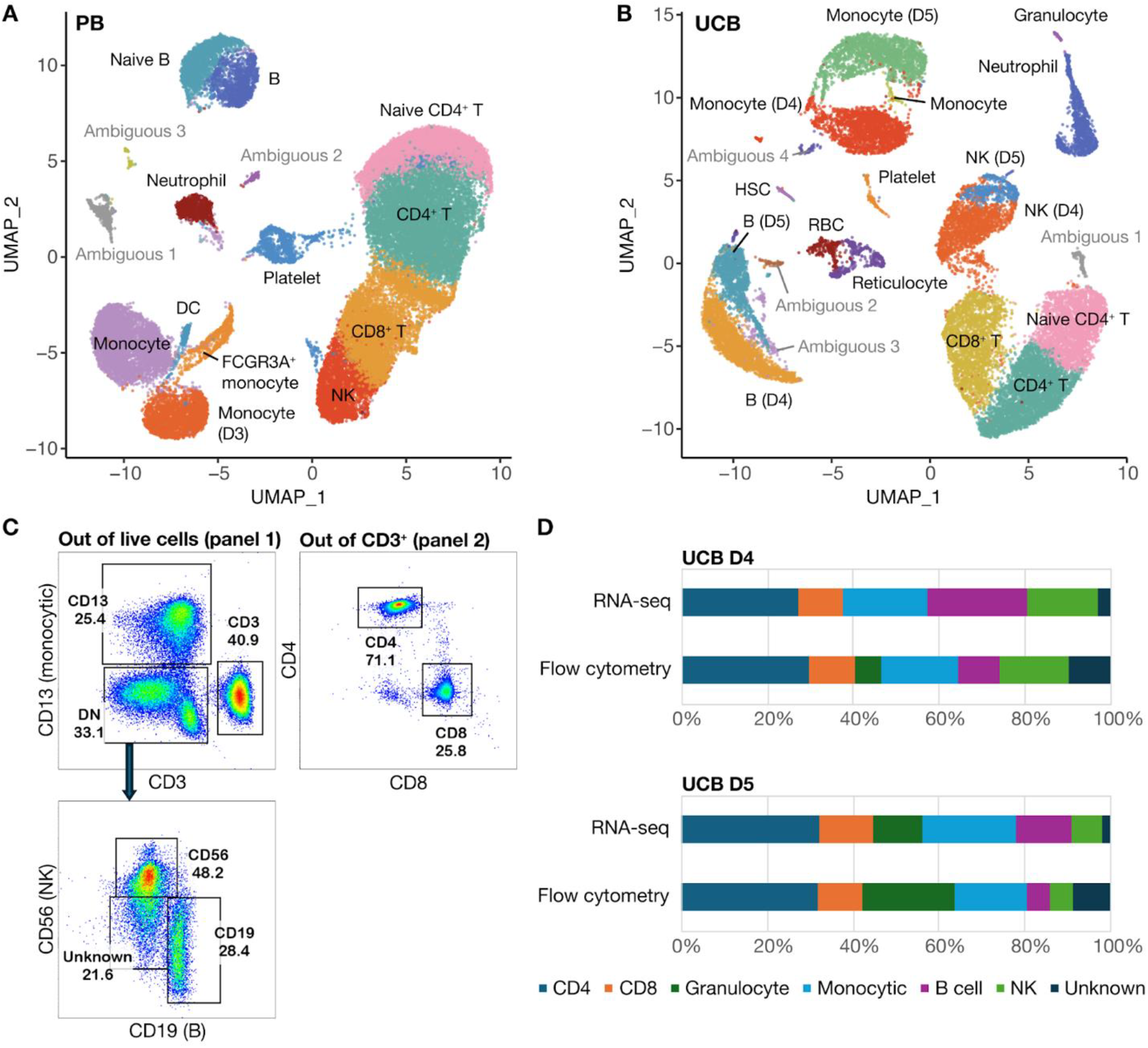
: scRNA-seq data with cells clustered according to donor type. (A) UMAP of clustered PB donor cells (n=3) (B) UMAP of clustered UCB donor cells (n=2) (C) Gating strategy for cell types identified by flow analysis (D) Comparative cell composition from flow cytometry versus scRNA-seq data.

## Discussion

The overall UCB MNC composition out of live cells consisted of T- and B-lymphocytes, natural killer (NK) cells, monocytes and granulocytes (Fig 2). UCB had a significantly higher percentage of CD19+ B-lymphocytes compared to adult PB (p=0.0125). There were no significant differences between the other cell populations. Based on range, UCB T-lymphocytes and granulocytes were the most heterogeneous, with the greatest inter-donor variability. We observed that most UCB donors had a high number of CD66b+ granulocytes compared to PB donors (Fig 2E). While elevated plasma G-CSF and neutrophil counts in UCB (25) have been previously reported, we were surprised to observe the presence of granulocytes since we were studying the MNC fraction, which should exclude polymorphonuclear cells. While Low-density Granulocytes (LDGs) found in the PB fraction have been reported in systemic lupus erythematosus (SLE) and other systemic autoimmune and autoinflammatory diseases (26), their role has not been studied extensively in healthy individuals or UCB. While we saw that most UCB donors had more granulocytes (Fig 2E), the trend was not significant due to a few outlier PB donors which could have underlying, inflammatory conditions.

UCB CD3+ T-lymphocytes were further characterized by CD4+ and CD8+ immunophenotyping, with a higher number of CD4+ subtypes (Fig 3). UCB had a higher percentage of CD3+CD4+ T-cells compared to PB (p=0.0265). Immaturity was determined by CD3+CD45RA+, detecting naive T-cell populations, and further by CD3+CD45RA+CD31+, detecting Recent Thymic Emigrants (RTE). Both subtypes were significantly higher amongst UCB samples compared to PB samples for both CD3+CD4+ and CD3+CD8+ populations, confirming the relatively immature properties of T-lymphocytes in human UCB. This specific characteristic of UCB might be associated with its lower incidence of graft-versus-host disease (GVHD) in the clinical setting, compared to bone marrow stem cell transplantation (11). Treg and TR1 CD3+CD4+ composition was low for UCB samples, suggesting that neonatal tolerance is largely mediated by the maternal immune system in keeping with the literature (34-36).

CD19+ B-lymphocytes were composed of a higher number of naive or transitional B cells versus memory B cells (Fig 4) (19). There was a significantly higher percentage of CD19+CD27+IgD-in PB compared to UCB, and a significantly higher percentage of CD19+IgD+CD27-in UCB compared to PB, again confirming the relatively immature properties of human UCB. Other studies have shown that although UCB-derived B cells have a more naive phenotype, they have a distinct transcriptional program conferring accelerated responsiveness to stimulation and facilitated IgA class switching (20). Mature NK cell composition was not significantly different between UCB and PB.

To our knowledge, this study of the immune composition of 50 UCB donors is the largest to date. The findings from flow cytometry were further supported by scRNA-seq data which showed concordance between flow cytometry and scRNA-seq data. While cells clustered into expected hematopoietic lineages observed in the literature (31, 32, 33), our results highlight that cells from PB donors tended to cluster together while cells from UCB donors tended to cluster separately (Fig 5C). There are separate clusters for UCB B cells, naive T cells, and Monocytes. In the PB dataset, 17 out of 18 clusters contained cells from multiple donors while in the UCB dataset, more clusters contained cells from an individual donor (6 out of 19 clusters). This suggests that while there may be variation at birth, cell types tend to converge in adulthood as the immune system matures. Another surprising observation is that naïve CD3 T cells from UCB cluster together regardless of whether they are CD4 or CD8 (Fig 5A). Our flow data support that the overwhelming majority of CD3 cells from UCB donors are naive: 87.79% of CD4 T cells and 94.87% of CD8 T cells are CD45RA+. These UCB T cells cluster separately from adult T cells, which in contrast form more differentiated clusters of CD4 naïve, CD4, and CD8 T cells. This adds to our understanding of adaptive immune maturation where T-Cell Receptor (TCR) stimulation in different immune and tissue contexts causes more differentiated but stereotyped effector cell types to emerge.

There are important considerations when interpreting the data from our study. First, our UCB samples were obtained from pregnant individuals at Lucile Packard Children’s Hospital (LPCH), so donor samples reflect the diverse ethnic backgrounds found in the Bay Area. LPCH serves insured, uninsured and underinsured families, and the patient population is largely Hispanic (56%), followed by Pacific Islander (13%), African American (11%), Asian (8%), Caucasian (7%) and unspecified (5%) (22). Additionally, we obtained the samples through the Binns Program for Cord Blood Research and the Stanford Blood Center, so conditions such as autoimmunity and neoplasms were screened and the patients consented were generally healthy (10). These factors influence the generalizability of our data. Notably, the health screening process might not have included specific conditions. Because our samples are de-identified, we cannot analyze certain specific findings, or make correlations of specific cell characteristics with specific medical histories.

One limitation of the study is the inability to capture all immune subsets. Since our aim was to study the MNC fraction from donor samples, polymorphonuclear cells in the densest fraction pelleted after Ficoll centrifugation were excluded. While we observed LDGs in the MNC fraction, the differences between PB and UCB were not statistically significant and may be less robust because we know less about the health status of the PB donors. These cells, the majority of which are Neutrophils, are sensitive to slight changes in temperature or ion concentrations (28), and could have been activated and lost during flow analysis or library preparation for RNA-seq. Notably, the Granulocyte population was present in the flow data of UCB donor 4 but absent from the scRNA-seq dataset (Table 2). While well-characterized in autoimmune conditions, the LDG population warrants further study in UCB and healthy individuals.

Human UCB is an overlooked resource and a material that is often discarded after delivery but may hold therapeutic potential. L. Bużańska et al. (2002) used the CD34-negative fraction of human UCB to obtain neural-like stem cells, demonstrating the high self-renewal potency of these populations (9). UCB derived regulatory T cells (Tregs) have also been investigated clinically as a source of cells for adoptive Treg transfer to prevent GVHD (16). Upon isolation, they have been shown to have comparable potency to adult peripheral blood (PB) derived Tregs (17) and to contain a higher proportion of naïve cells (CD45RA+), which have longer-term phenotypic and epigenetic stability as compared to memory Tregs (18). We were able to study this cell source in-depth through the Binns Program for Cord Blood Research. Our study provides a robust characterization of the immune subset composition of a large cohort of human umbilical cord blood and control adult peripheral blood donors through flow cytometry immunophenotyping. We were able to combine this with finer grained scRNA-seq analysis which showed concordance in the proportion of immune cell types but captured more differentiated subtypes of cells. These analyses yield the intriguing possibility that immune heterogeneity of individuals at birth gives way to more stereotyped immune subsets as the immune system is exposed to the external environment and undergoes maturation.

## Acknowledgments

Binns Program for Cord Blood Research

We acknowledge Maurizio Morri from the Chan Zuckerberg Biohub Network as well as Jennifer Cory and Ginger Exley from the Stanford Center for Definitive and Curative Medicine for their assistance and coordination for this project.

## References

1. Osgood EE, Riddle MC, Mathews TJ. Aplastic anemia treated with daily transfusions and intravenous marrow; case report. Ann Intern Med. 1939;13:357–67. doi: 10.7326/0003-4819-13-2-357

2. Henig I, Zuckerman T. Hematopoietic stem cell transplantation-50 years of evolution and future perspectives. Rambam Maimonides Med J. 2014;5(4):e0028. Published 2014 Oct 29. doi:10.5041/RMMJ.10162

3. Newcomb JD, Sanberg PR, Klasko SK, Willing AE. Umbilical cord blood research: current and future perspectives. Cell Transplant. 2007;16(2):151–158.

4. Gluckman E, Rocha V. History of the clinical use of umbilical cord blood hematopoietic cells. Cytotherapy. 2005;7(3):219–227. doi:10.1080/14653240510027136

5. Paloczi K. Immunophenotypic and functional characterization of human umbilical cord blood mononuclear cells. Leukemia. 1999;13 Suppl 1:S87–S89. doi:10.1038/sj.leu.2401318

6. D’Arena G, Musto P, Cascavilla N, et al. Flow cytometric characterization of human umbilical cord blood lymphocytes: immunophenotypic features. Haematologica. 1998;83(3):197–203.

7. Aldenhoven M, Kurtzberg J. Cord blood is the optimal graft source for the treatment of pediatric patients with lysosomal storage diseases: clinical outcomes and future directions. Cytotherapy. 2015;17(6):765–774. doi:10.1016/j.jcyt.2015.03.609

8. Rocha V, Chastang C, Souillet G, et al. Related cord blood transplants: the Eurocord experience from 78 transplants. Eurocord Transplant group. Bone Marrow Transplant. 1998;21 Suppl 3:S59–S62.

9. Buzańska L, Machaj EK, Zabłocka B, Pojda Z, Domańska-Janik K. Human cord blood-derived cells attain neuronal and glial features in vitro. J Cell Sci. 2002;115(Pt 10):2131–2138. doi:10.1242/jcs.115.10.2131

10. Mantri S, Sheikali A, Binns C, et al. The Binns Program for Cord Blood Research: A novel model of cord blood banking for academic biomedical research. Placenta. 2021;103:50–52. doi:10.1016/j.placenta.2020.10.018

11. Rocha V, Wagner JE Jr, Sobocinski KA, et al. Graft-versus-host disease in children who have received a cord-blood or bone marrow transplant from an HLA-identical sibling. Eurocord and International Bone Marrow Transplant Registry Working Committee on Alternative Donor and Stem Cell Sources. N Engl J Med. 2000;342(25):1846–1854. doi:10.1056/NEJM200006223422501

12. Mantri S, Reinisch A, Dejene BT, et al. CD34 expression does not correlate with immunophenotypic stem cell or progenitor content in human cord blood products. Blood Adv. 2020;4(21):5357–5361. doi:10.1182/bloodadvances.2020002891

13. Wynn L, Wilson MG, Leonforte C. Manufacturing of CD34 + HPC-enriched, high-purity mononuclear cell products from umbilical cord blood [published online ahead of print, 2023 Feb 3]. Cell Tissue Bank. 2023;10.1007/s10561-023-10070-8. doi:10.1007/s10561-023-10070-8

14. Amand M, Iserentant G, Poli A, et al. Human CD56dimCD16dim Cells As an Individualized Natural Killer Cell Subset. Front Immunol. 2017;8:699. Published 2017 Jun 19. doi:10.3389/fimmu.2017.00699

15. Keever CA. Characterization of cord blood lymphocyte subpopulations. J Hematother. 1993;2(2):203–206. doi:10.1089/scd.1.1993.2.203

16. Claudio G. Brunstein, Jeffrey S. Miller, Qing Cao, David H. McKenna, Keli L. Hippen, Julie Curtsinger, Todd DeFor, Bruce L. Levine, Carl H. June, Pablo Rubinstein, Philip B. McGlave, Bruce R. Blazar, John E. Wagner; Infusion of ex vivo expanded T regulatory cells in adults transplanted with umbilical cord blood: safety profile and detection kinetics. Blood 2011; 117 (3): 1061–1070. doi: 10.1182/blood-2010-07-293795

17. Claudio G. Brunstein, Jeffrey S. Miller, David H. McKenna, Keli L. Hippen, Todd E. DeFor, Darin Sumstad, Julie Curtsinger, Michael R. Verneris, Margaret L. MacMillan, Bruce L. Levine, James L. Riley, Carl H. June, Chap Le, Daniel J. Weisdorf, Philip B. McGlave, Bruce R. Blazar, John E. Wagner; Umbilical cord blood–derived T regulatory cells to prevent GVHD: kinetics, toxicity profile, and clinical effect. Blood 2016; 127 (8): 1044–1051. doi: 10.1182/blood-2015-06-653667

18. Petra Hoffmann, Ruediger Eder, Tina J. Boeld, Kristina Doser, Biserka Piseshka, Reinhard Andreesen, Matthias Edinger, Only the CD45RA+ subpopulation of CD4+CD25high T cells gives rise to homogeneous regulatory T-cell lines upon in vitro expansion, Blood, Volume 108, Issue 13, 2006, Pages 4260–4267, ISSN 0006-4971, 10.1182/blood-2006-06-027409.

19. Sanz I, Wei C, Jenks SA, Cashman KS, Tipton C, Woodruff MC, Hom J, Lee FE. Challenges and Opportunities for Consistent Classification of Human B Cell and Plasma Cell Populations. Front Immunol. 2019 Oct 18;10:2458. doi: 10.3389/fimmu.2019.02458. PMID: 31681331; PMCID: PMC6813733.

20. Bettina Budeus, Artur Kibler, Martina Brauser, Ekaterina Homp, Kevin Bronischewski, J. Alexander Ross, Andre Görgens, Marc A. Weniger, Josefine Dunst, Taras Kreslavsky, Symone Vitoriano da Conceição Castro, Florian Murke, Christopher C. Oakes, Peter Rusch, Dimitrios Andrikos, Peter Kern, Angela Köninger, Monika Lindemann, Patricia Johansson, Wiebke Hansen, Anna-Carin Lundell, Anna Rudin, Jan Dürig, Bernd Giebel, Daniel Hoffmann, Ralf Küppers, Marc Seifert; Human Cord Blood B Cells Differ from the Adult Counterpart by Conserved Ig Repertoires and Accelerated Response Dynamics. J Immunol 15 June 2021; 206 (12): 2839–2851. 10.4049/jimmunol.2100113

21. Yi Zhao, Xiao Li, Weihua Zhao, Jingwan Wang, Jiawei Yu, Ziyun Wan, Kai Gao, Gang Yi, Xie Wang, Bingbing Fan, Qinkai Wu, Bangwei Chen, Feng Xie, Jinghua Wu, Wei Zhang, Fang Chen, Huanming Yang, Jian Wang, Xun Xu, Bin Li, Shiping Liu, Yong Hou, Xiao Liu, Single-cell transcriptomic landscape of nucleated cells in umbilical cord blood, GigaScience, Volume 8, Issue 5, May 2019, giz047, 10.1093/gigascience/giz047

22. Stanford Pediatrics Residency. (n.d.). Educational settings. https://med.stanford.edu/peds/prospective-applicants/educational-settings.html

23. Sen P, Kemppainen E, Orešič M. Perspectives on Systems Modeling of Human Peripheral Blood Mononuclear Cells. Front Mol Biosci. 2018 Jan 9;4:96. doi: 10.3389/fmolb.2017.00096. PMID: 29376056; PMCID: PMC5767226.

24. Luo OJ, Lei W, Zhu G, Ren Z, Xu Y, Xiao C, Zhang H, Cai J, Luo Z, Gao L, Su J, Tang L, Guo W, Su H, Zhang ZJ, Fang EF, Ruan Y, Leng SX, Ju Z, Lou H, Gao J, Peng N, Chen J, Bao Z, Liu F, Chen G. Multidimensional single-cell analysis of human peripheral blood reveals characteristic features of the immune system landscape in aging and frailty. Nat Aging. 2022 Apr;2(4):348–364. doi: 10.1038/s43587-022-00198-9. Epub 2022 Apr 18. PMID: 37117750.

25. Laver J, Duncan E, Abboud M, Gasparetto C, Sahdev I, Warren D, Bussel J, Auld P, O’Reilly RJ, Moore MA. High levels of granulocyte and granulocyte-macrophage colony-stimulating factors in cord blood of normal full-term neonates. J Pediatr. 1990 Apr;116(4):627–32. doi: 10.1016/s0022-3476(05)81617-8. PMID: 1690796.

26. Carmona-Rivera C, Kaplan MJ. Low-density granulocytes in systemic autoimmunity and autoinflammation. Immunol Rev. 2023 Mar;314(1):313–325. doi: 10.1111/imr.13161. Epub 2022 Oct 28. PMID: 36305174; PMCID: PMC10050110.

27. Hao Y, Stuart T, Kowalski MH, Choudhary S, Hoffman P, Hartman A, Srivastava A, Molla G, Madad S, Fernandez-Granda C, Satija R. Dictionary learning for integrative, multimodal and scalable single-cell analysis. Nat Biotechnol. 2024 Feb;42(2):293–304. doi: 10.1038/s41587-023-01767-y. Epub 2023 May 25. PMID: 37231261; PMCID: PMC10928517.

28. Cui C, Schoenfelt KQ, Becker KM, Becker L. Isolation of polymorphonuclear neutrophils and monocytes from a single sample of human peripheral blood. STAR Protoc. 2021 Sep 21;2(4):100845. doi: 10.1016/j.xpro.2021.100845. PMID: 34604813; PMCID: PMC8473578.

29. Jain V, Yang WH, Wu J, Roback JD, Gregory SG, Chi JT. Single Cell RNA-Seq Analysis of Human Red Cells. Front Physiol. 2022 Apr 20;13:828700. doi: 10.3389/fphys.2022.828700. PMID: 35514346; PMCID: PMC9065680.

30. Kapellos TS, Bonaguro L, Gemünd I, Reusch N, Saglam A, Hinkley ER, Schultze JL. Human Monocyte Subsets and Phenotypes in Major Chronic Inflammatory Diseases. Front Immunol. 2019 Aug 30;10:2035. doi: 10.3389/fimmu.2019.02035. PMID: 31543877; PMCID: PMC6728754.

31. Zhao Y, Li X, Zhao W, Wang J, Yu J, Wan Z, Gao K, Yi G, Wang X, Fan B, Wu Q, Chen B, Xie F, Wu J, Zhang W, Chen F, Yang H, Wang J, Xu X, Li B, Liu S, Hou Y, Liu X. Single-cell transcriptomic landscape of nucleated cells in umbilical cord blood. Gigascience. 2019 May 1;8(5):giz047. doi: 10.1093/gigascience/giz047. PMID: 31049560; PMCID: PMC6497034.

32. Shi X, Ma W, Duan S, Shi Q, Wu S, Hao S, Dong G, Li J, Song Y, Liu C, Lin X, Yuan Y, Deng Q, Xu J, Bai S, Hou Y, Liu C, Liu L. Single-cell transcriptional diversity of neonatal umbilical cord blood immune cells reveals neonatal immune tolerance. Biochem Biophys Res Commun. 2022 Jun 11;608:14–22. doi: 10.1016/j.bbrc.2022.03.132. Epub 2022 Mar 28. PMID: 35381424.

33. Bunis DG, Bronevetsky Y, Krow-Lucal E, Bhakta NR, Kim CC, Nerella S, Jones N, Mendoza VF, Bryson YJ, Gern JE, Rutishauser RL, Ye CJ, Sirota M, McCune JM, Burt TD. Single-Cell Mapping of Progressive Fetal-to-Adult Transition in Human Naive T Cells. Cell Rep. 2021 Jan 5;34(1):108573. doi: 10.1016/j.celrep.2020.108573. PMID: 33406429; PMCID: PMC10263444.

34. Salvany-Celades M, van der Zwan A, Benner M, Setrajcic-Dragos V, Bougleux Gomes HA, Iyer V, Norwitz ER, Strominger JL, Tilburgs T. Three Types of Functional Regulatory T Cells Control T Cell Responses at the Human Maternal-Fetal Interface. Cell Rep. 2019 May 28;27(9):2537-2547.e5. doi: 10.1016/j.celrep.2019.04.109. PMID: 31141680.

35. Erlebacher A. Immunology of the maternal-fetal interface. Annu Rev Immunol. 2013;31:387–411. doi: 10.1146/annurev-immunol-032712-100003. Epub 2013 Jan 3. PMID: 23298207.

36. Tilburgs T, Roelen DL, van der Mast BJ, de Groot-Swings GM, Kleijburg C, Scherjon SA, Claas FH. Evidence for a selective migration of fetus-specific CD4+CD25bright regulatory T cells from the peripheral blood to the decidua in human pregnancy. J Immunol. 2008 Apr 15;180(8):5737–45. doi: 10.4049/jimmunol.180.8.5737. PMID: 18390759.

