## Supplemental Figures for "Immune Composition of the Mononuclear Cell Fraction of Human Umbilical Cord Blood"

### Supplemental Tables

**Supplemental Table 1.** Panel design (Beckman Coulter CytoFLEX Flow Cytometer). (A) Initial UCB and aPB samples were processed with Original Panels. Later revision prompted changes to Panels 1 and 2 and addition of Panel 3 for a more comprehensive analysis of cell types. (B) Clone and catalog number for antibodies used.

1A

| MARKER | FLUOROPHORE |
|---|---|
| Live/Dead | Ghost Dye BV510 |
| CD3 | PE-CF594 |
| CD13 | PE |
| CD14 | BV421 |
| CD19 | BV605 |
| CD56 | PE-Cy7 |
| HLA-DR | FITC |
| CD27 | APC-Cy7 |
| CD16 | APC |

Original Panel 1 (General)

| MARKER | FLUOROPHORE |
|---|---|
| Live/Dead | Ghost Dye BV510 |
| CD3 | PE-CF594 |
| CD4 | BV421 |
| CD8 | BV605 |
| CD25 | APC-Cy7 |
| CD127 | FITC |
| CD45RA | APC |
| CD31 | PE-Cy7 |

Original Panel 2 (T Cell)

| MARKER | FLUOROPHORE |
|---|---|
| CD3 | PE-CF594 |
| CD13 | PE |
| CD14 | BV421 |
| CD19 | BV605 |
| CD56 | PE-Cy7 |
| HLA-DR | FITC |
| CD66b | APC-Cy7 |
| CD16 | APC |
| Live-Dead | Ghost Dye BV510 |

Updated Panel 1 (General)

| MARKER | FLUOROPHORE |
|---|---|
| CD3 | PE-CF594 |
| CD4 | BV421 |
| CD8 | BV605 |
| CD31 | PE-Cy7 |
| CD45RA | APC |
| CD19 | BV650 |
| IgD | FITC |
| CD27 | APC-Cy7 |
| Live-Dead | Ghost Dye BV510 |
| CD14 | BV510 (dump) |

Updated Panel 2 (T & B Lymphocytes)

| MARKER | FLUOROPHORE |
|---|---|
| CD3 | PE-CF594 |
| CD4 | APC Cy7 |
| CD25 | BV605 |
| CD127 | BV421 |
| CD49b | FITC |
| LAG3 | APC |
| Live-Dead | Ghost Dye BV510 |
| CD14 | BV510 (dump) |

Panel 3 (Regulatory T Lymphocytes)

1B

| MARKER | FLUOROPHORE | CLONE | CATALOG NUMBER |
|---|---|---|---|
| Live/Dead | Ghost Dye BV510 | - | 50-201-4118 |
| CD3 | PE-CF594 | UCHT1 | BDB562280 |
| CD13 | PE | WM15 | BDB560998 |
| CD14 | BV421 | MφP9 | BDB563744 |
| CD19 | BV605 | SJ25C1 | BDB562654 |
| CD56 | PE-Cy7 | B159 | BDB557747 |
| HLA-DR | FITC | LN3 | 50-112-2195 |
| CD27 | APC-Cy7 | O323 | 50-165-906 |
| CD16 | APC | 3G8 | BDB561248 |
| CD4 | BV421 | SK3 | BDB566907 |
| CD8 | BV605 | SK1 | BDB564115 |
| CD25 | APC-Cy7 | M-A251 | BDB561782 |
| CD127 | FITC | HIL-7R-M21 | BDB560549 |
| CD45RA | APC | 5H9 | BDB561210 |
| CD31 | PE-Cy7 | WM59 | BDB563651 |
| CD66b | APC-Cy7 | G10F5 | 305125 |
| IgD | FITC | IA6-2 | 50-213-1878 |
| CD14 | BV510 (dump) | MφP9 | BDB563079 |
| CD4 | APC Cy7 | RPA-T4 | BDB557871 |
| CD25 | BV605 | BC96 | BDB567572 |
| CD127 | BV421 | hIL-7R-M21 | BDB562436 |
| CD49b | FITC | REA188 | 130-100-335 |
| LAG3 | APC | REA351 | 130-119-567 |

**Supplemental Table 2**. Detailed statistics for MNC and PBMC comparisons.

<table>
<tr><th>Pop</th><th>N Obs</th><th>Variable</th><th>Mean</th><th>Std Dev</th><th>Lower 95%</th><th>Upper 95%</th><th>N</th><th>t Value</th><th>Pr > |t|</th></tr>
<tr><th></th><th></th><th></th><th></th><th></th><th>CL for Mean</th><th>CL for Mean</th><th></th><th></th><th></th></tr>
<tr><td>CD3_live</td><td>49</td><td>UCB</td><td>42.888</td><td>10.369</td><td>39.91</td><td>45.867</td><td>49</td><td>28.95</td><td><.0001</td></tr>
<tr><td></td><td></td><td>PB</td><td>42.368</td><td>20.727</td><td>33.178</td><td>51.558</td><td>22</td><td>9.59</td><td><.0001</td></tr>
<tr><td>CD19_live</td><td>49</td><td>UCB</td><td>8.07</td><td>3.168</td><td>7.16</td><td>8.98</td><td>49</td><td>17.83</td><td><.0001</td></tr>
<tr><td></td><td></td><td>PB</td><td>5.836</td><td>3.181</td><td>4.426</td><td>7.247</td><td>22</td><td>8.61</td><td><.0001</td></tr>
<tr><td>CD56_live</td><td>49</td><td>UCB</td><td>10.573</td><td>5.439</td><td>9.01</td><td>12.135</td><td>49</td><td>13.61</td><td><.0001</td></tr>
<tr><td></td><td></td><td>PB</td><td>12.95</td><td>8.883</td><td>9.012</td><td>16.888</td><td>22</td><td>6.84</td><td><.0001</td></tr>
<tr><td>Monocytes_live</td><td>49</td><td>UCB</td><td>19.852</td><td>6.375</td><td>18.021</td><td>21.683</td><td>49</td><td>21.8</td><td><.0001</td></tr>
<tr><td></td><td></td><td>PB</td><td>21.386</td><td>11.676</td><td>16.071</td><td>26.701</td><td>21</td><td>8.39</td><td><.0001</td></tr>
<tr><td>Granulocytes_live.</td><td>49</td><td>UCB</td><td>15.402</td><td>12.667</td><td>11.763</td><td>19.04</td><td>49</td><td>8.51</td><td><.0001</td></tr>
<tr><td></td><td></td><td>PB</td><td>12.386</td><td>20.668</td><td>3.223</td><td>21.55</td><td>22</td><td>2.81</td><td>0.0105</td></tr>
<tr><td>CD4_CD3</td><td>50</td><td>UCB</td><td>71.953</td><td>6.029</td><td>70.24</td><td>73.667</td><td>50</td><td>84.39</td><td><.0001</td></tr>
<tr><td></td><td></td><td>PB</td><td>65.132</td><td>12.749</td><td>59.479</td><td>70.784</td><td>22</td><td>23.96</td><td><.0001</td></tr>
<tr><td>CD8_CD3</td><td>50</td><td>UCB</td><td>25.331</td><td>6.04</td><td>23.615</td><td>27.048</td><td>50</td><td>29.65</td><td><.0001</td></tr>
<tr><td></td><td></td><td>PBMC</td><td>25.445</td><td>10.182</td><td>20.931</td><td>29.96</td><td>22</td><td>11.72</td><td><.0001</td></tr>
</table>

| | | | | | | | | | |
|---|---|---|---|---|---|---|---|---|---|
| CD45RA_CD4. | 50 | UCB | 87.794 | 13.354 | 83.999 | 91.589 | 50 | 46.49 | <.0001 |
| | | PB | 42.373 | 11.784 | 37.148 | 47.597 | 22 | 16.87 | <.0001 |
| CD45RA_CD8 | 50 | UCB | 94.869 | 6.522 | 93.015 | 96.722 | 50 | 102.85 | <.0001 |
| | | PB | 61.4 | 19.731 | 52.652 | 70.148 | 22 | 14.6 | <.0001 |
| CD31_Naive CD4. | 50 | UCB | 79.671 | 12.141 | 76.22 | 83.122 | 50 | 46.4 | <.0001 |
| | | PB | 56.164 | 20.084 | 47.259 | 65.068 | 22 | 13.12 | <.0001 |
| CD31_Naive CD8 | 50 | UCB | 99.529 | 0.555 | 99.371 | 99.686 | 50 | 1268.63 | <.0001 |
| | | PB | 84.227 | 16.175 | 77.056 | 91.399 | 22 | 24.42 | <.0001 |
| Treg_CD4 | 46 | UCB | 3.706 | 1.69 | 3.204 | 4.208 | 46 | 14.87 | <.0001 |
| | | PBMC | 3.139 | 1.758 | 2.264 | 4.013 | 18 | 7.57 | <.0001 |
| Tr1_CD4 | 20 | UCB | 0.269 | 0.376 | 0.093 | 0.445 | 20 | 3.2 | 0.0047 |
| | | PB | 0.47 | 0.583 | 0.053 | 0.887 | 10 | 2.55 | 0.0313 |
| Memory B _ CD19 | 24 | UCB | 3.189 | 2.229 | 2.248 | 4.13 | 24 | 7.01 | <.0001 |
| | | PB | 24.486 | 8.223 | 19.738 | 29.234 | 14 | 11.14 | <.0001 |
| Naive B _ CD19 | 24 | In | 80.121 | 10.402 | 75.729 | 84.514 | 24 | 37.74 | <.0001 |
| | | PB | 53.186 | 13.714 | 45.268 | 61.104 | 14 | 14.51 | <.0001 |
| CD16_CD56. | 49 | UCB | 73.319 | 15.553 | 68.851 | 77.786 | 49 | 33 | <.0001 |

| | | | | | | | | | |
|---|---|---|---|---|---|---|---|---|---|
| | | PB | 79.886 | 13.84 | 73.75 | 86.023 | 22 | 27.07 | <.0001 |
| HLA-DR- _ GM | 49 | UCB | 33.292 | 23.187 | 26.632 | 39.952 | 49 | 10.05 | <.0001 |
| | | PB | 22.482 | 29.014 | 9.618 | 35.346 | 22 | 3.63 | 0.0016 |
| HLA-DR+ _ GM | 49 | UCB | 61.114 | 21.952 | 54.808 | 67.419 | 49 | 19.49 | <.0001 |
| | | PB | 71.255 | 28.839 | 58.468 | 84.041 | 22 | 11.59 | <.0001 |
| CD66b+ _ HLADR- | 24 | UCB | 91.707 | 20.216 | 83.17 | 100.243 | 24 | 22.22 | <.0001 |
| | | PB | 98.092 | 1.851 | 96.974 | 99.211 | 13 | 191.09 | <.0001 |
| CD66b+ _ HLADR+ | 24 | UCB | 0.658 | 0.71 | 0.358 | 0.958 | 24 | 4.54 | 0.0001 |
| | | PB | 0.264 | 0.348 | 0.063 | 0.465 | 14 | 2.84 | 0.0138 |

**Supplemental Table 3**. Mean differences ± SEM of PBMC and UCB populations. Differences in mean values are calculated as PBMC-UCB means. NS: p-value not significant, *: p-value<0.05, **: p-value:<0.01 ***: p-value=<0.001 ****: p-value<0.0001.

| Pop | Difference Between Means (PB-UCB) ± SEM | 95% Confidence Interval | P-Value | P-Value Summary | Significantly different (P < 0.05)? |
|---|---|---|---|---|---|
| CD3/live | -0.4598 ± 4.654 | -10.03 to 9.109 | 0.9221 | ns | No |
| CD19/live | -2.125 ± 0.8129 | -3.767 to -0.4826 | 0.0125 | * | Yes |
| CD56/live | 2.525 ± 2.030 | -1.629 to 6.679 | 0.2236 | ns | No |
| Monocytes/live | 1.516 ± 2.596 | -3.809 to 6.842 | 0.5640 | ns | No |

| | | | | | |
|---|---|---|---|---|---|
| Granulocytes/live | -2.904 ± 4.757 | -12.64 to 6.835 | 0.5465 | ns | No |
| CD4/CD3 | -6.729 ± 2.854 | -12.61 to -0.8528 | 0.0265 | * | Yes |
| CD8/CD3 | 0.2231 ± 2.331 | -4.554 to 5.000 | 0.9244 | ns | No |
| CD45RA/CD4 | -45.36 ± 3.143 | -51.69 to -39.03 | <0.0001 | **** | Yes |
| CD45RA/CD8 | -33.39 ± 4.313 | -42.31 to -24.47 | <0.0001 | **** | Yes |
| CD31/Naive CD4 | -23.39 ± 4.622 | -32.86 to -13.92 | <0.0001 | **** | Yes |
| CD31/Naive CD8 | -15.23 ± 3.450 | -22.40 to -8.051 | 0.0002 | *** | Yes |
| Treg/CD4 | -0.5132 ± 0.4834 | -1.500 to 0.4739 | 0.2968 | ns | No |
| Tr1/CD4 | 0.2631 ± 0.1958 | -0.1591 to 0.6854 | 0.2016 | ns | No |
| Memory B / CD19 | 21.35 ± 2.245 | 16.54 to 26.16 | <0.0001 | **** | Yes |
| Naive B / CD19 | -26.88 ± 4.236 | -35.67 to -18.10 | <0.0001 | **** | Yes |
| CD16/CD56 | 6.617 ± 3.695 | -0.8247 to 14.06 | 0.0800 | ns | No |
| HLA-DR+ / GM | 10.21 ± 6.896 | -3.826 to 24.26 | 0.1482 | ns | No |
| HLA-DR- / GM | -10.64 ± 7.038 | -24.95 to 3.670 | 0.1399 | ns | No |
| CD66b+ / HLADR+ | -0.3311 ± 0.1731 | -0.6824 to 0.02016 | 0.0639 | ns | No |
| CD66b+ / HLADR- | 6.578 ± 4.156 | -2.006 to 15.16 | 0.1267 | ns | No |

**Supplemental Table 4.** Pre- and post-quality control (QC) cell estimates for each donor.

| Donors | Estimated Cells (Cell Ranger) | Cell Number Post QC |
|---|---|---|
| PB D1 | 11734 | 10912 |
| PB D2 | 12944 | 12036 |
| PB D3 | 15022 | 13967 |
| UCB D4 | 11877 | 10939 |
| UCB D5 | 11635 | 10789 |
| UCB D6 | 111851 | NA |

**Supplemental Table 5.** Cell types and their lineage specifying genes.

| | **Cell Type** | **Genes** | **Naive** |
|---|---|---|---|
| 1 | B cell | MS4A1 | IGHD |
| 2 | CD4 | CD3D, CD3E, CD4 | SELL, CCR7 |

| | | | |
|---|---|---|---|
| 3 | CD8 | CD3D, CD3E, CD8A, CD8B | SELL, CCR7 |
| 4 | Neutrophil / Granulocyte | AQP9, LIMK2 | |
| 5 | HSC | CD34 | |
| 6 | RBC / Reticulocyte | HBA1, HBA2 | |
| 7 | Platelets | PPBP, GP1BB | |
| 8 | Monocytic | LYZ, HLA-DRA, S100A8, CD86 | |
| 9 | Non-classical / intermediate Monocytes | FCGR3A (CD16) | |
| 10 | DC | FCER1A | |
| 11 | NK | GNLY | |

### Supplemental Figures

**1A**

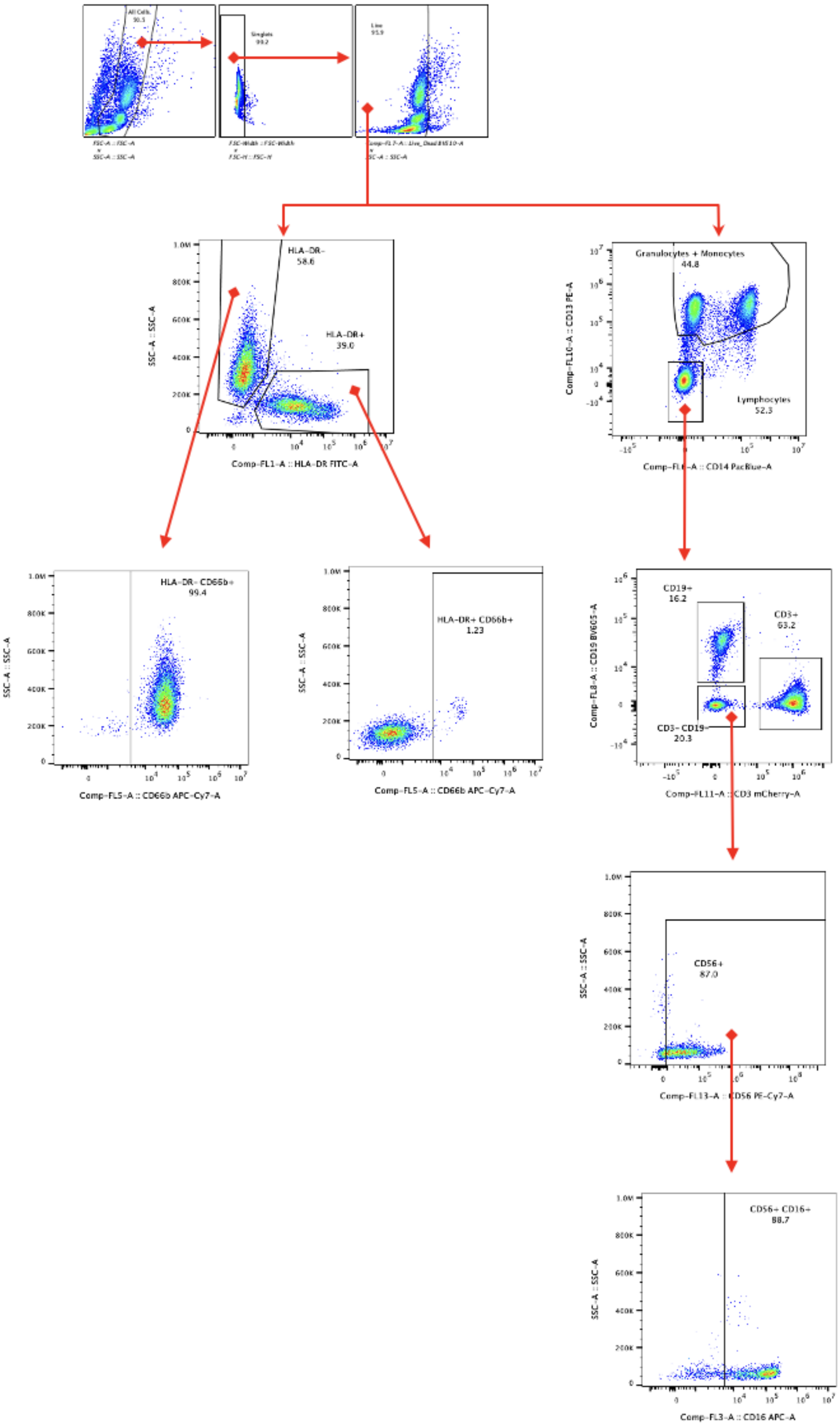
HLA-DR-
58.6
HLA-DR+
39.0
SSC-A :: SSC-A
Comp-FL1-A :: HLA-DR FITC-A
Granulocytes + Monocytes
44.8
Lymphocytes
52.3
Comp-FL10-A :: CD13 PE-A
CD14 PacBlue-A
HLA-DR- CD66b+
99.4
Comp-FL5-A :: CD66b APC-Cy7-A
HLA-DR+ CD66b+
1.23
Comp-FL5-A :: CD66b APC-Cy7-A
CD19+
16.2
CD3+
63.2
CD3- CD19-
20.3
Comp-FL8-A :: CD19 BV605-A
Comp-FL11-A :: CD3 mCherry-A
CD56+
87.0
Comp-FL13-A :: CD56 PE-Cy7-A
CD56+ CD16+
88.7
Comp-FL3-A :: CD16 APC-A

**1B**

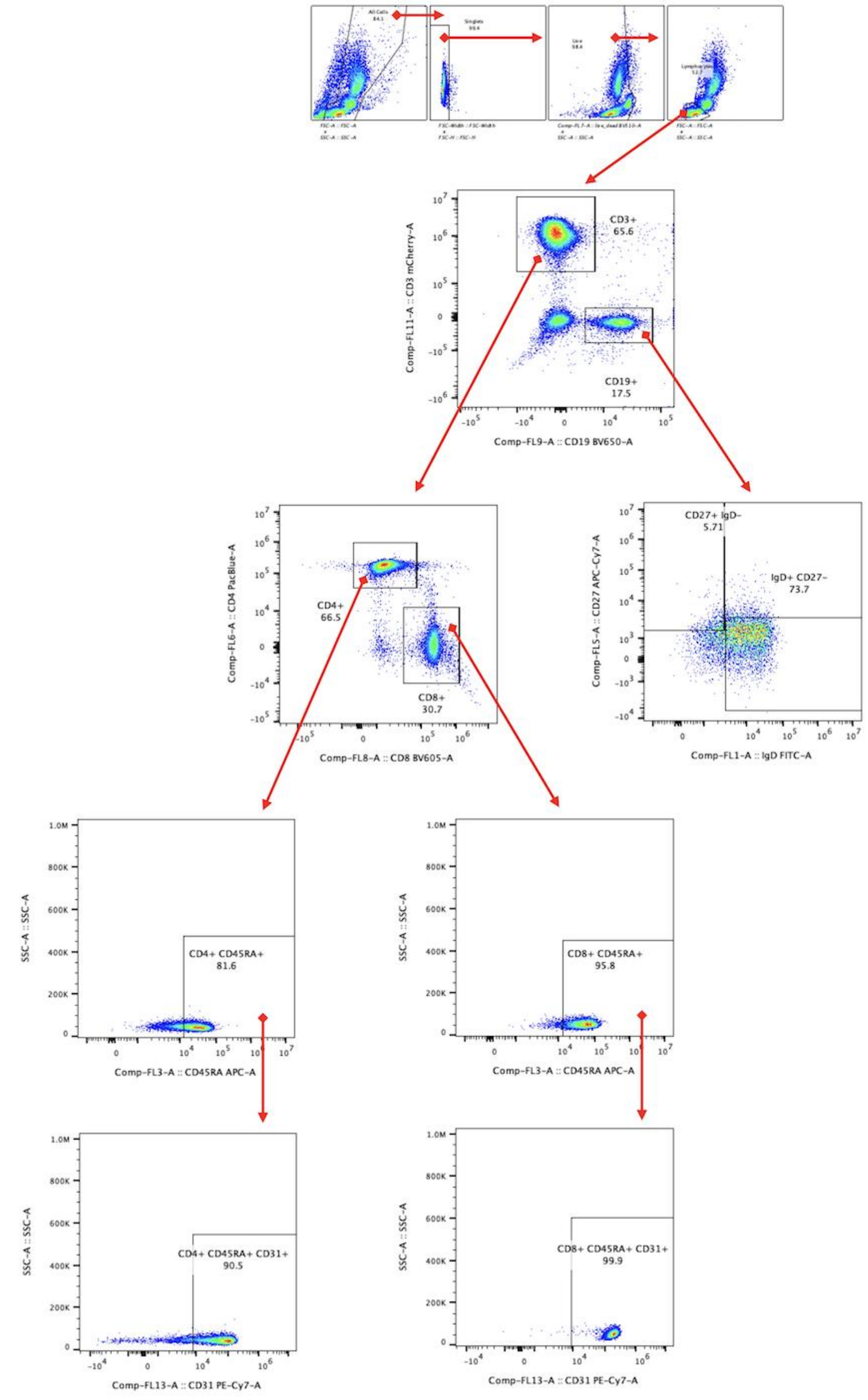

CD3+
65.6
CD19+
17.5
Comp-FL11-A :: CD3 mCherry-A
Comp-FL9-A :: CD19 BV650-A
CD4+
66.5
CD8+
30.7
Comp-FL6-A :: CD4 PacBlue-A
Comp-FL8-A :: CD8 BV605-A
CD27+ IgD-
5.71
IgD+ CD27-
73.7
Comp-FL5-A :: CD27 APC-Cy7-A
Comp-FL1-A :: IgD FITC-A
CD4+ CD45RA+
81.6
CD8+ CD45RA+
95.8
SSC-A :: SSC-A
Comp-FL3-A :: CD45RA APC-A
CD4+ CD45RA+ CD31+
90.5
CD8+ CD45RA+ CD31+
99.9
Comp-FL13-A :: CD31 PE-Cy7-A


**1C**

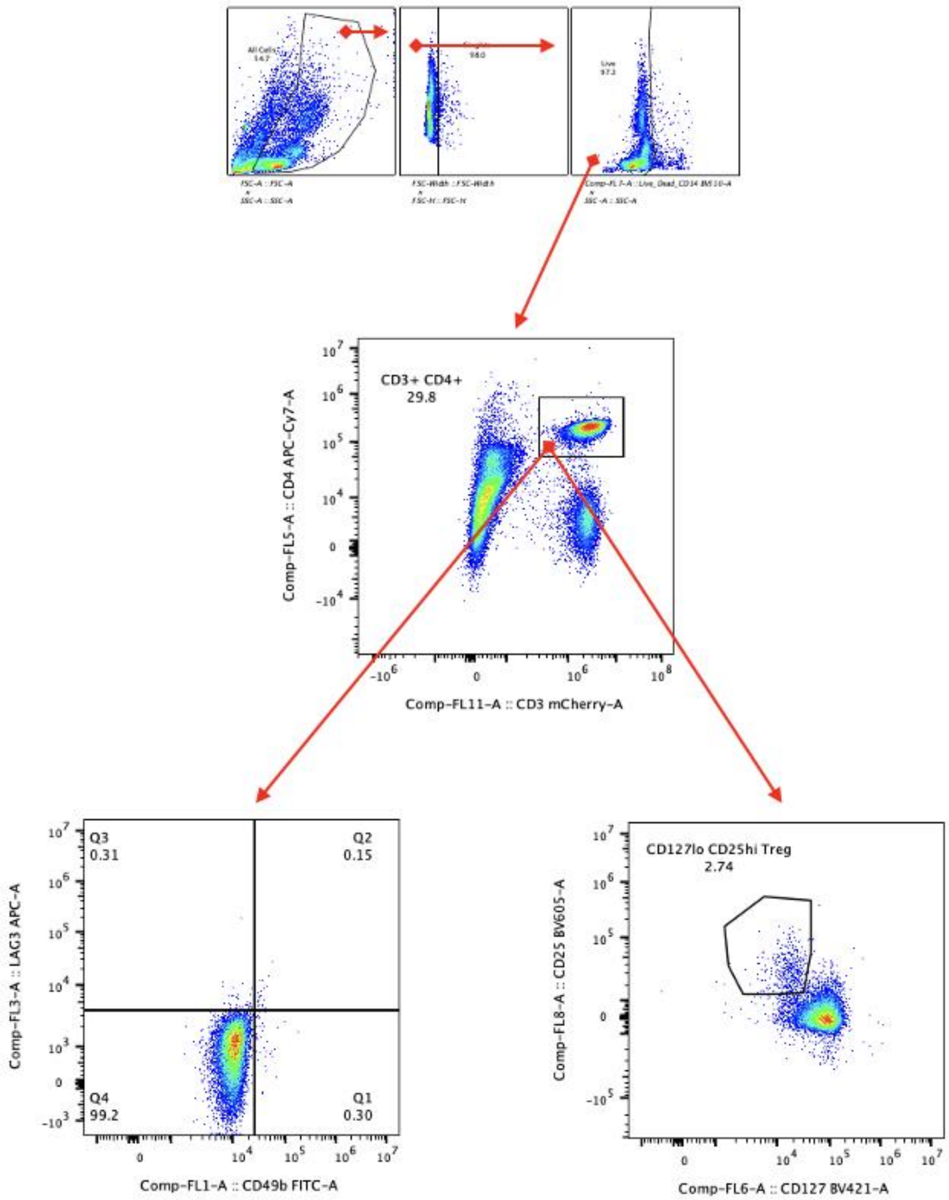


**Supplemental Figure 1.** Gating strategy for (1A) Panel 1, (1B) Panel 2, and (1C) Panel 3.

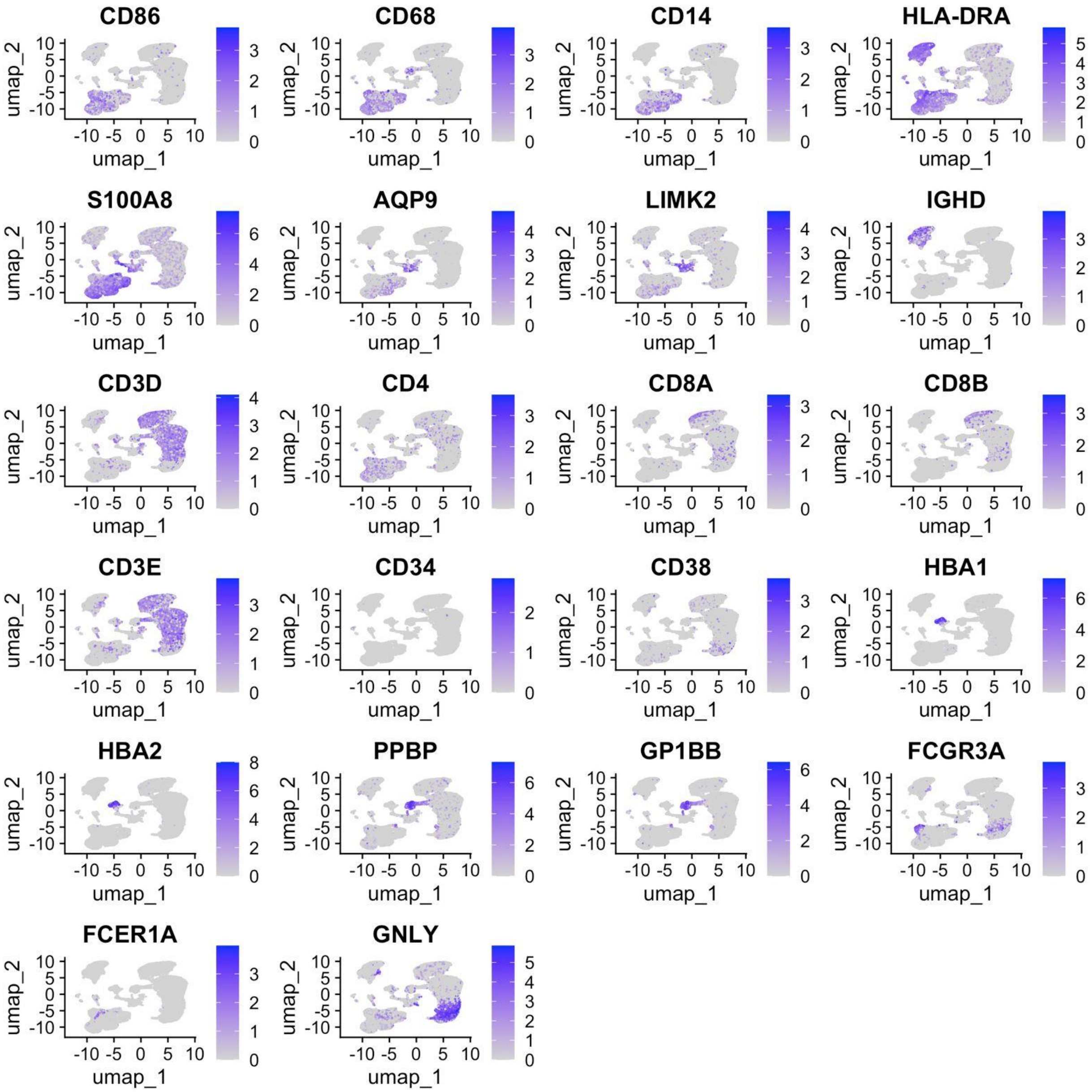


**Supplemental Figure 2**. UMAPs of the combined PB and UCB dataset depicting gene expression profiles of key lineage-specifying genes. PB: n=3 donors, UCB: n=2 donors.

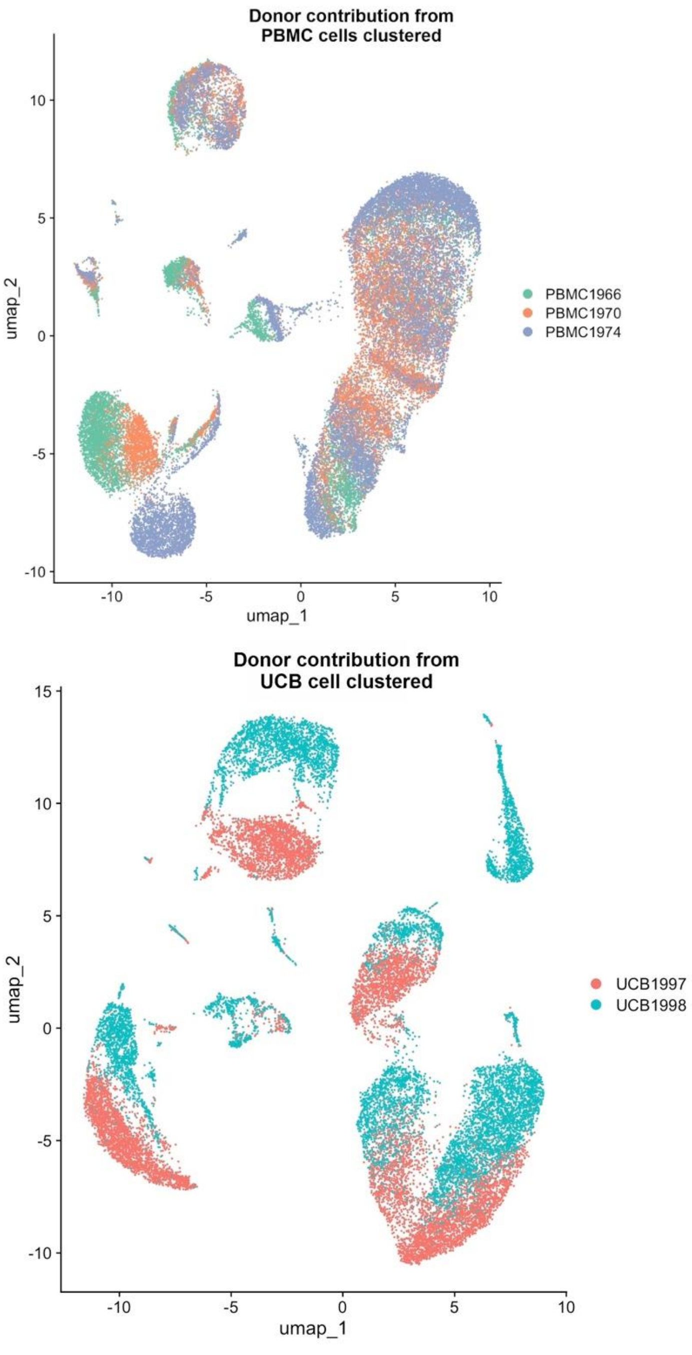


**Supplemental Figure 3.** UMAPs of clustered UCB (left) and PB (right) cells, coloured by donor. PB: n=3 donors, UCB: n=2 donors.
